# Triggered functional dynamics of AsLOV2 by time-resolved electron paramagnetic resonance at high magnetic fields

**DOI:** 10.1101/2022.10.12.511365

**Authors:** Shiny Maity, Brad D. Price, C. Blake Wilson, Arnab Mukherjee, Matthieu Starck, David Parker, Maxwell Z. Wilson, Janet E. Lovett, Songi Han, Mark S. Sherwin

## Abstract

We present time-resolved Gd-Gd electron paramagnetic resonance (TiGGER) at 240 GHz for tracking inter-residue distances during a protein’s mechanical cycle in the solution state. TiGGER makes use of Gd-sTPATCN as spin labels, whose favorable qualities include a spin-7/2 EPR-active center, short linker, narrow intrinsic linewidth, and virtually no anisotropy at high fields (8.6 T) when compared to nitroxide spin labels. Using TiGGER, we determined that upon light activation, the C-terminus and N-terminus of AsLOV2 separate in less than 1 s and relax back to equilibrium with a time constant of approximately 60 s. TiGGER revealed that the light-activated long-range mechanical motion is slowed in the Q513A variant of AsLOV2 and is correlated to the similarly slowed relaxation of the optically excited chromophore as described in recent literature. TiGGER has the potential to valuably complement existing methods for the study of triggered functional dynamics in proteins.

## 1 Introduction

Proteins are fundamental building blocks of life. Understanding their function is key to understanding biological processes; this desire to understand has culminated in more than 190,000 structures being logged into the Protein Data Bank.[1] The thoroughness with which static structures have been mapped leads one to begin considering a functional “movie” – observing multiple biologically relevant amino acid sites move in real time – which can be combined to create a 3D rendition of site-specific motion. However, most state-of-the-art structural biology tools require the protein to be immobilized (cryo-electron microscopy, solid-state NMR, or double electron electron resonance – DEER) or mechanically inhibited (X-ray crystallography) which may cause a significant amount of information to be lost (*e.g*., time-dependence, environmental effects, pH effects).[2, 3] By rapidly freezing an ensemble of proteins after triggering a conformational change, several structural biology tools have been used to elucidate the triggered functional dynamics of proteins [4–6]. However, ideally, a “movie” would be “filmed” in real-time, by *in vitro* tracking of the positions of residues at or near physiological temperatures in the solution state.

Pioneering work of Steinhoff and Hubbell demonstrated that continuous-wave electron paramagnetic resonance (cwEPR) at X-band (9.5 GHz, 0.35 T) can be used to report on site-specific structural changes in proteins upon activation under physiological conditions.[7–9] Such experiments made use of nitroxide-based labels with anisotropic *g* and hyperfine tensors that provide extensive knowledge about the local environment in which the spin label resides. Specifically, the label tumbling rate reveals the rigidity and the local spatial constraints of the spin labeled residue in 3D space. When triggered mechanical activation alters the local rigidity and environment, the cwEPR lineshape can pick up on these changes to report on protein conformation. Unfortunately, the sensitivity of nitroxide-based cwEPR lineshapes to their local environment makes it challenging to extract distances from the small contribution of dipolar broadening to spectra of proteins doubly labeled with nitroxide molecules.[10, 11] Pulsed electron dipolar spectroscopies such as DEER solve this problem by isolating the dipolar interaction from other factors, and hence are widely used for measuring distances between protein residues,[12–18] but, with a few exceptions (*e.g*., [19]), are always performed at cryogenic temperatures.

In this paper, we show that cwEPR lineshape measurements at high magnetic fields (8.6 T) and frequencies (240 GHz), combined with the recently-developed Gd(III) spin label Gd-sTPATCN [20] (“Gd-NO_3_Pic” in [21]), enable time-resolved measurement of distance changes between labeled residues of a photoresponsive protein in solution at room temperature. Unlike nitroxide spin labels, the spin-7/2 paramagnetic center in Gd(III) is shielded from its local environment, and has a nearly isotropic g-tensor with negligible hyperfine interactions. These attributeslead to an extremely narrow cwEPR line associated with the −1/2 to 1/2 central transition Gd spin labels at high magnetic fields.[22] The lineshape of this central transition thus reports sensitively on the distance to nearby Gd spin labels, even at room temperature, as long as the dipolar coupling is strong enough for molecular tumbling to not completely average out its orientation-dependent interaction.[23]. We call our method time-resolved gadolinium-gadolinium electron paramagnetic resonance (TiGGER).

We apply TiGGER to a protein in the Light, Oxygen, and Voltage (LOV) family, which undergo reversible structural changes in response to stimulation.[24–32] In fact, these structural changes are central to the utility of LOV proteins as engineered optogenetic actuators to establish light-dependent mechanical control over various aspects of cellular function.[27, 33–35] However, the bioengineering of efcient LOV protein-based actuators is done empirically by systematic mutation and screening.[27, 29, 30] Clearly, the design of efcient actuators would benefit from an in-depth understanding of site-specific and inter-site movements *in vitro*, which cannot be adequately resolved with currently available biophysical techniques.

Specifically, we study AsLOV2, a phototropin 1 LOV2 domain from *Avena sativa* (oats)[35–37]. Upon light activation, a flavin mononucleotide (FMN) within the chromophore binding pocket forms an adduct with an internal cysteine that initiates a cascade of conformational changes, culminating in the unfolding of a peripheral J*α*-helix. There have been numerous EPR studies that characterize the effects of mutations on the light-sensitive chromophore binding pocket [24–26, 28, 38–40], as well as reports on light-induced flavin [32] and structural changes (at cryogenic temperatures) [31]. However, none focused on directly measuring the transient, time-resolved, long-range structural changes that LOV proteins undergo to execute their optogenetic functions. The mutation of a conserved Q513 glutamine residue is thought to slow down LOV protein activation, but the nature of the coupling between light activation and structural changes in LOV is a subject of active debate [3, 41–44]. Hence, for TIGGER’s debut, we present time-resolved measurements of the J*α*-helix unfolding in WT and the Q513A mutant of AsLOV2.

## 2 Results

We select for labeling a residue on the J*α*-helix (near the C-terminus, 537) that is expected to unfold and move significantly upon light activation, as well as a second site on the N-terminus (406) that is expected to remain relatively still.[1, 47] The two sites are mutated to cysteine residues (mutations T406C and E537C) and subsequently spin-labeled with a common nitroxide based label, MTSL, or Gd-sTPATCN (labeling procedure in S.I. 1.3). Numerical simulations on Multiscale Modeling of Macromolecules software (version 2018.2) show that when crystallized, nitroxide-based MTSL labels attached to residues 406 and 537 are ~2.63 nm apart; this distance in the solution state is not known for certain, but we expect it to remain similar (±0.4 nm).[48, 49] Upon light activation in the solution state, it is expected that sites 406 and 537 become more separated and that TiGGER is sensitive to the distance change induced by their motion.[47, 50]

The photoresponse of spin labeled AsLOV2 was first established using UV-Vis spectroscopy. As shown in Fig. 1 (middle left), the spectrum changed most dramatically at 447 nm while under blue (450 nm) light illumination due to changes in the local environment of the FMN chromophore. Time-dependent UV-Vis measurements (middle right) clearly show that the protein is photoswitching and relaxing as a result of this illumination, and that site directed mutagenesis and spin labeling (SDSL) at sites 406 and 537 does not affect the photoresponse of AsLOV2 (see S.I. Figs. 5-8).

**Figure 1:**
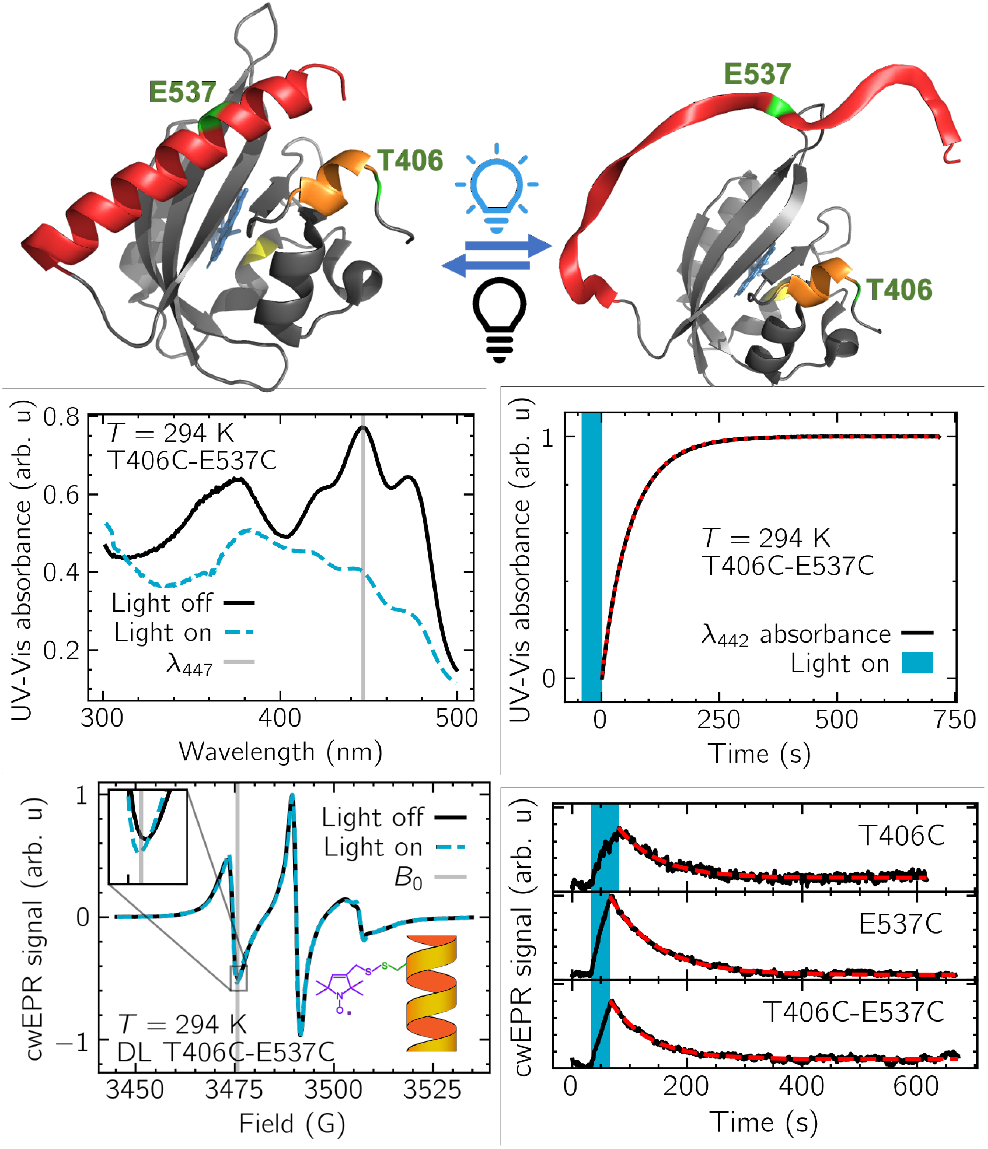
Photoresponse of AsLOV2. (Top) PYMOL(v2.5.2)-generated structure demonstrating AsLOV2 structural change that occurs after 450 nm illumination (left: dark-state, PDB 2V1A; right: hypothesized lit-state).[45, 46] The residues 537 and 406 that were labeled in this paper are marked green on J*α*-helix (C-terminus, shown in red) and A-*α*-helix (N-terminus, shown in orange), respectively. (Middle left) UV-Vis absorption spectra of AsLOV2 T406C-E537C with (dashed blue line) and without (solid black line) blue light activation (Thorlabs, Inc. LIU470A). The vertical gray line indicates the wavelength at which the lifetime of the protein was measured. (Middle right) The lifetime of the protein (*τ* = 65.06 ± 0.03 s) after activation with blue light was measured by recording the UV-Vis absorbance at 447 nm. (Bottom left) X-band cwEPR spectra of AsLOV2 labeled at the residues 537 and 406 with MTSL, a nitroxide-based standard EPR spin label. (Bottom right) Transient X-band EPR demonstrating similar signals from doubly and singly MTSL-labeled AsLOV2. Time constants for the fits (dashed red lines) were *τ*_*T*406*C*_ = 70.1 ± 1.3 s, *τ*_*E*537*C*_ = 80.5 ± 0.6 s, *τ*_*T*406*C−E*537*C*_ = 66.2 ± 0.8 s. Transient data and their fits showed no significant change in amplitude after light activation between singly (SL) and doubly labeled (DL) samples. Field values used for time-dependent measurements were done at the position of maximum change near 3475 G, within ±1 G.

To confirm that the protein actuates, and that the actuation is correlated to its photoresponse, we recorded X-band EPR spectra of proteins labeled with the nitroxide spin label MTSL. The X-band EPR spectra showed changes between the light on and off states, which corresponded to a change in MTSL’s local environment (see Fig. 1, bottom left). Time-resolved X-band EPR experiments showed similar (65 s < *τ* < 81 s) relaxation times to those for photoactivation obtained from UV-Vis spectra (*τ* = 65.06 ± 0.3 s), confirming that the protein is moving upon light activation (Fig. 1 bottom right). There was no distinguishable difference between singly and doubly MTSL-labeled samples, which implies that the changes seen in the cwEPR spectral amplitude under light activation are not due to changes in the inter-spin label couplings, but instead to changes in the local environment of the individual MTSL spin labels. Such changes are expected, as the J-*α* and A-*α* helices unfolding can readily alter the tumbling rate, anisotropy, and, potentially, tertiary contact of the MTSL spin label. This observation illustrates the need for spin labels sensitive to intra-protein distance-dependent dipolar coupling.

Engineered gadolinium(III)-sTPATCN labels offer the opportunity for dipolar-sensitive EPR at the distance ranges relevant to AsLOV2 mechanical action (~1-4 nm [23]). Gd-sTPATCN has many qualities that make it favorable for TiGGER: it is centrosymmetric, high-spin (*S* = 7/2), has a short, rigid linker, and negligible hyperfine coupling.[20] Additionally, its central (*m*_*s*_ = −1/2 → 1/2) transition is extremely narrow at high fields (~5 G at 8.6 T) because of its extremely small (Δ*ω* ≈ 2*π* * 485 MHz) zero-field splitting (ZFS).[21] To first order, ZFS does not contribute to the central transition linewidth, and to second order, linewidth scales as 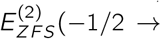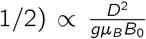, which is inversely proportional to *B*_0_.[20, 51] Because of the extremely narrow central transition, and because dipolar magnetic fields from states with both |*m*_*s*_ | = 1/2 and |*m*_*s*_| > 1/2 contribute to dipolar broadening of Gd(III), spin-spin distances approaching 4 nm broaden the cwEPR linewidth at 8.6 T.[23] The favorable high-spin and narrow linewidth properties of Gd labels have been extensively utilized for DEER measurements under cryogenic conditions.[52–59]

It is important for TiGGER that the sample contain a minimal fraction of singly-labeled protein. To remove singly labeled proteins, an already spin-labeled preparation of double mutants was further incubated with biotin-maleimde (BM) and filtered with a streptavidin-agarose (SA) resin, which removed any proteins with free cysteines (*i.e*., imperfectly labeled, see Fig. 2, top and right, and S.I. 1.4 for filtering details).

**Figure 2:**
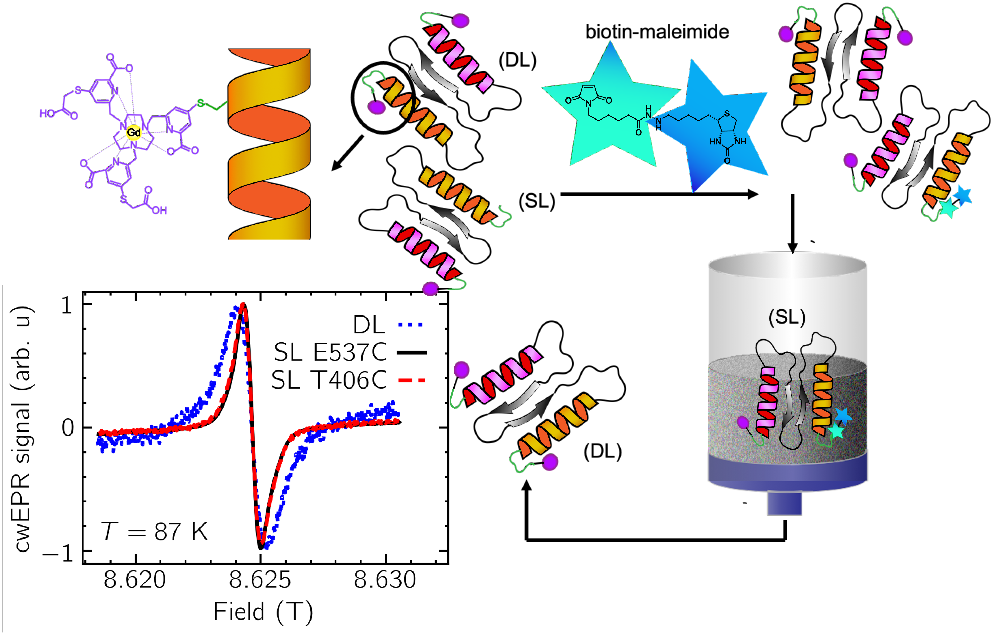
Site-directed spin labeling of AsLOV2 with Gd-sTPATCN. (Top and right) Enrichment of DL AsLOV2. The reaction products from the spin-labeling are mixed with biotin maleimide. Singly-labeled AsLOV2 with free cysteine is trapped in the column with streptavidin agarose and the fully labeled DL T406C-E537C gets through the column. (Bottom left) Comparison of 240 GHz cwEPR lineshapes of AsLOV2 singly and doubly labeled with Gd-sTPATCN. Lineshapes are normalized to highlight dipolar broadening. The experiment was done at 87 K to eliminate effects of motional averaging. Dotted blue line, solid black line, and dashed red line correspond to AsLOV2 samples labeled at residues 406-437, 537, and 406, respectively.

To ensure that we are able to observe dipolar broadening of doubly labeled (DL) AsLOV2, we first performed 240 GHz cwEPR experiments at 87 K that froze out tumbling and enhanced dipolar broadening (spectrometer details outlined in S.I. 1.8). Dipolar broadening was observed and is shown in Fig. 2 (bottom left): the doubly Gd-labeled AsLOV2 EPR line has a larger peak-to-peak linewidth, as well as broader wings, than that of singly labeled (SL) AsLOV2 at sites 406 or 537. The lineshapes of the two singly labeled proteins are essentially indistinguishable, emphasizing the insensitivity of this spin label to its local environment.

The principal result of this paper is shown in Fig. 3. The top row shows that, at room temperature, the cwEPR spectra of singly labeled proteins at either site 406 or 537 do not change upon light activation; while the protein does activate (see Fig. 1, middle and bottom right), the Gd-sTPATCN label is not sensitive to it. DL AsLOV2 samples (Fig. 1, bottom left), by comparison, show a significant lineshape narrowing upon light activation at room temperature. We attribute this change to a reduction in dipolar broadening between spin labels at sites 406 and 537, as these sites moved apart when the J-*α* and A-*α* helices unfolded. This result is in contrast to the X-band EPR experiment presented in Fig. 1 (bottom row): the lineshape change observed at X-band did not depend on whether the sample was doubly or singly labeled. Therefore, with TiGGER, we were able to directly observe a change in dipolar coupling – and therefore protein motion – after activation by an external stimulus in solution state. TiGGER of doubly Gd-sTPATCN labeled AsLOV2 yielded a time constant for relaxation to equilibrium similar to those from X-band EPR and UV-Vis spectroscopy results (*τ*_*TiGGER*_ = 51.9 ± 0.3 s, 65 s < *τ*_*X–band*_ < 81 s, *τ*_*UV–V is*_ = 65.06 ± 0.03 s). This observation confirms that mechanical relaxation is correlated to that of the local environment of MTSL and the triplet state of FMN.

**Figure 3:**
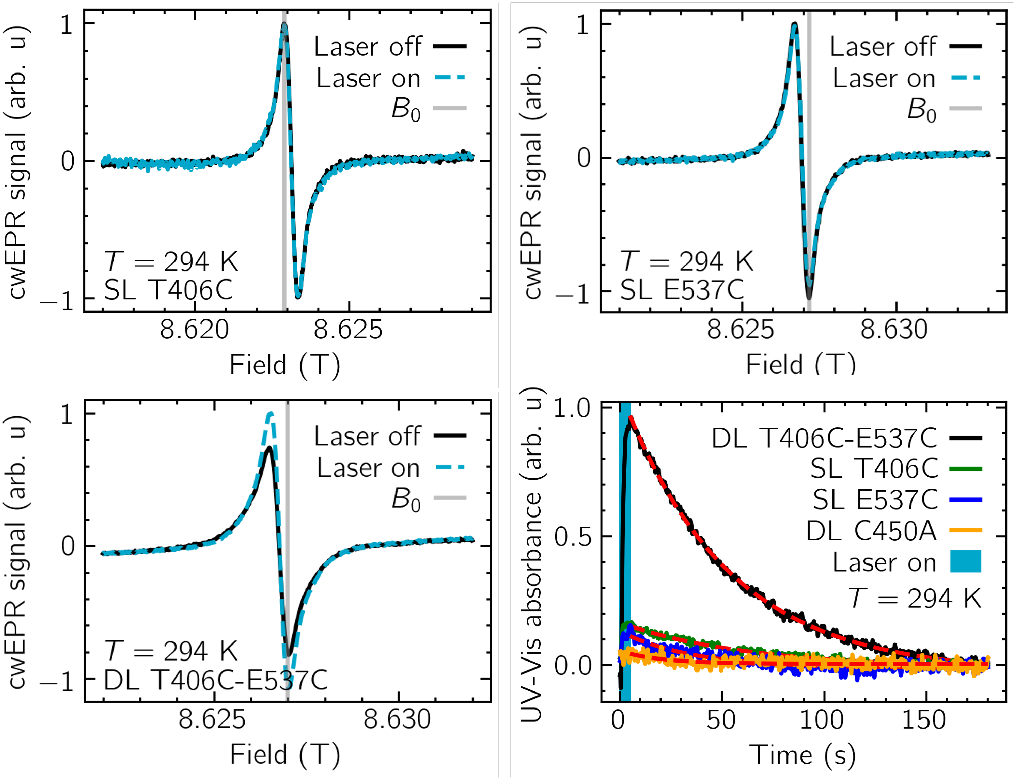
Effect of laser illumination on cwEPR spectra of Gd-labeled AsLOV2. SL cwEPR spectra of AsLOV2 residues 406 (top left) and 537 (top right) demonstrating that the spectrum with the laser off is unchanged when the laser is turned on. (Bottom left) DL (sites 406-537) cwEPR spectra of AsLOV2 demonstrating that the spectrum with the laser off is narrowed when the laser is turned on. Laser off spectra are shown in solid black and laser on spectra are shown in dashed blue. *B*_0_, where maximum time-dependent change occurred, is shown on all three plots by a vertical gray line. Note that *B*_0_ is not the same for all three samples; the field value with maximum change was chosen for each time-dependent experiment. (Bottom right) cwEPR time-dependent signal change of SL and DL Gd-AsLOV2 due to laser illumination at *T* = 294 K, shown by solid blue line (solid black, green, blue, and orange lines correspond to DL T406C-E537C, SL T406C, SL E537C, and DL T406C-E537C C450A, respectively). Overlaid best fits (dashed red lines) of the exponentials provide time constants of *τ*_T406C-E537C_ = 51.9 ± 0.3 s, *τ*_T406C_ = 62.5 ± 1.8 s, *τ*_E537C_ = 34.6 ± 1.9 s, and *τ*_T406C-E537C C450A_ = 20.8 ± 3.4 s. All plots are normalized to the magnitude of DL T406C-E537C signal change. See S.I. 2.4 hypothesized cause of nonzero SL change.

The internal cysteine residue C450 is intimately coupled to the chromophore. Previous studies have shown that mutation of this conserved residue leads to complete inhibition of the photocycle and suppression of the associated secondary structural changes.[35, 60] As a negative control for TiGGER, experiments were completed on DL AsLOV2 with the internal cysteine mutated to alanine (C450A). No illumination-dependent effects were observed under static or transient EPR conditions (static shown in S.I. Fig. 10; transient shown in orange in Fig. 3, bottom right). Our UV-Vis experiments on C450A DL AsLOV2 corroborated this result and confirmed that laser illumination initiated no spectroscopic changes, consistent with previously published results. Therefore, we can conclude that the line narrowing seen by TiGGER in DL AsLOV2 is due to a light-activated inter-label distance increase caused by photoactivation.

We next turn our attention to a topic of current interest in the literature: conserved glutamine residue (Q513).[3, 41, 47, 50, 61–63] Mutation to an uncharged, non-polar amino acid (*e.g*., alanine), is known to slow down photoactivation and relaxation of AsLOV2. TiGGER is well equipped to inform us on whether a Q513A mutation also alters the long-range structural changes as manifested in the J*α*-helix unfolding. The Q513 site is located on the I*α*-sheet, interacts with the J*α*-helix, and is in the immediate proximity of the chromophore binding site. It is suspected that, upon light activation, this residue switches its hydrogen bonding pattern with the chromophore and plays a key role in the transmission of stress to the J*α*-helix, which causes it to unfold.[41, 43] In order to elucidate the consequence of Q513A mutation in the unfolding of the J*α*-helix, we generated a Q513A variant of DL AsLOV2 with Gd-sTPATCN spin label attached to the same Gd (III) sites (T406C and E537C) as before. We first confirmed, by transient UV-Vis, that the Q513A mutation does not inhibit photoswitching of DL AsLOV2, but does slow the photoactivity of the chromophore kinetics by a factor of approximately 2 (*τ*_Q513A DL_ = 135.4 ± 0.3, s see Fig. 4, right) compared to DL AsLOV2 (*τ*_DL_ = 65.06 ± 0.03 s). Further, measurement of distance changes via TiGGER between sites 406 and 537 via cwEPR showed that the Q513A mutation experienced similar, though appreciably less, dipolar narrowing to that of DL T406C-E537C upon 450 nm illumination. We also observed slowing of the TiGGER relaxation after activation (approx. 3x, *τ*_Q513A DL_ = 174.5 ± 0.5 s, see Fig. 4, right) that was comparable to what was observed by UV-Vis kinetics. This result is consistent with earlier studies on AsLOV2 using circular dichroism spectroscopy by other groups, where light activated changes were observed in molar ellipticity per residue, [*θ*]MRW, at 208 nm and 220 nm in the mutant Q513L AsLOV2.[3, 41, 47] Our results show that the Q513 residue plays an important role in modulating the structural changes of J*α*-helix upon blue light illumination but is not indispensible for motion, as the unfolding of the J*α*-helix is slowed but not suppressed.

**Figure 4:**
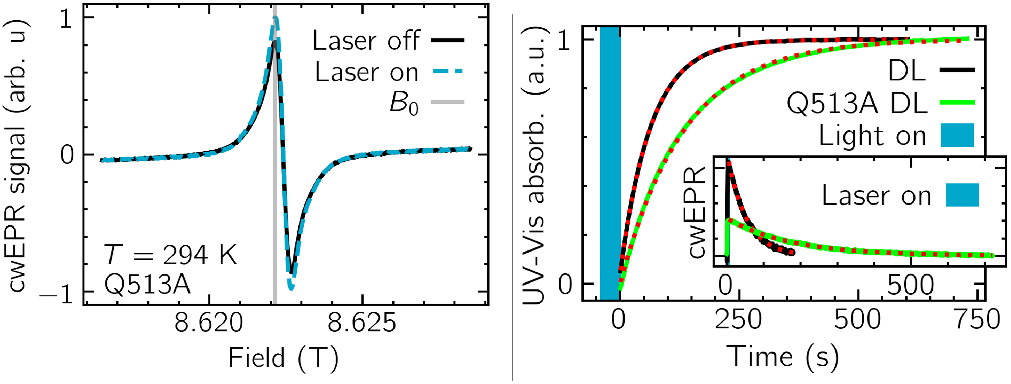
Q513A slowing of chromophore and mechanical photocycles as detected by TiGGER. (Left) cwEPR spectra of Q513A DL T406C-E537C AsLOV2 showing light-activated change between dark (solid black line) and lit (dashed blue line) states. (Right) Time-resolved UV-Vis demonstrates a slowing of the chromophore photocycle in AsLOV2 after illumination (vertical blue line). Black and green solid lines represent T406C-E537C at λ_447_ and Q513A DL T406C-E537C at λ_442_, respectively (see S.I. 1.6 for discussion on monitoring slightly different wavelengths). Best fits are shown by dashed red lines with respective time constants of *τ*_DL_ = 65.06 ± 0.03 s and *τ*_Q513A DL_ = 135.4 ± 0.3 s. (Inset) TiGGER at *B*_0_ with and without Q513A mutation demonstrating that Q513A slows mechanical motion. Best fits give *τ*_*DL*_ = 51.9 ± 0.3 s and *τ*_Q513A DL_ = 174.5 ± 0.5 s. Relative amplitudes are preserved after normalization.

## 3 Discussion

Upon light activation, the FMN chromophore is excited to a triplet state and reacts with the nearby C450 residue. It forms a covalent bond between the C(4a) atom of the chromophore and the sulfur atom of C450.[64] This cysteinyl adduct formation is accompanied by destabilization of the J*α*-helix in the C-terminus of the protein. The recovery rates of the covalent adduct to the dark state vary widely across LOV domains, on a timescale of seconds to days.[65–68] This widely varying photocycle is interesting; it is believed that the adduct decay is base-catalyzed, limited by a proton transfer step.[69] The presence of a glutamine residue, Q513, in hydrogen-bonding interaction with the chromophore, seems to be the only LOV residue in proximity to the chromophore that is capable of catalyzing this deprotonation process.[41] Our UV-Vis experiments on Q513A AsLOV2, along with the previous studies by others, show that mutation to the glutamine residue slows the photocycle, but does not prohibit it.[3, 43, 47] Studies in the literature have suggested that this could be potentially explained by effects of hydration of the LOV domain, where water molecules can directly act as a base catalyst by entering the chromophore binding pocket and hydrogen-bonding with the FMN chromophore.[3, 42, 66, 70, 71] Indeed, we observed that the chromophore and mechanical photocycles are still intact, even in the absence of the glutamine residue, albeit 2-3× slower. The mechanical slowing indicates that Q513 is important for catalysis near the FMN core, as well as for driving mechanical movement between the N and C-termini. However, the Q513 residue is not uniquely capable of driving J*α* unfolding, consistent with discussions in the literature [3, 41, 47].

Our observation that the J*α*-helix unfolds in the wild type, as well as Q513A, highlights the complementarity of TiGGER and other structural biology tools. Very recent high-resolution X-ray crystal structures of the mutant Q513L in ref. [3] show that related structural changes (sub-Å displacements) are induced by light in the Q513L mutant and the wild type. This result implies, however, that the magnitude of such changes cannot be readily seen by X-ray crystallography. It has long been known that when crystallized, the J*α*-helix remains folded upon light activation because unfolding is blocked by steric constraints [46]. In contrast, because TiGGER is performed in solution state, the J*α*-helix is free to unfold and readily does so. Solution-state NMR studies on Q513L AsLOV2 conducted by others show that, upon illumination, it is likely that the majority of the Q513L AsLOV2 population remains in the dark state. They report that this result is somewhat ambiguous, since chemical shift changes are sensitive to modifications in electronic structure of the FMN (upon adduct formation) and can obscure the effect of inter-site distance changes.[41] TiGGER, on the other hand, is not sensitive to FMN electron configurations (SL samples showed no light-activated change) and can unambiguously inform us about the effect of mutations on both unfolding yield and relaxation rate of the J*α*-helix. A reduction in photoactivity may also explain the reduced TiGGER amplitude observed for Q513A. By examining results from X-ray crystallography, solution-state NMR, and TiGGER together we arrive at a more complete view of triggered functional dynamics in AsLOV2.

To our knowledge, besides TiGGER, the only method capable of tracking triggered distance changes between a pair of specific residues on a protein in solution state is FRET. FRET relies on the distance-dependent non-radiative energy transfer from donor to acceptor fluorophores that have been introduced site-specifically into the macromolecule of interest.[72] Recent advances have also made it possible for its application on a single molecule level.[73–75] However, the application of FRET to explore dynamics in a light sensitive protein can be complicated due to the need for additional correction factors to resolve the spectral crosstalk between the light sensitive protein and donor-acceptor pairs.[76]

Future work for TiGGER on photoresponsive proteins will focus on enhancing the contrast between dark and lit states, mitigating the effects of rotational averaging, and making TiGGER a more quantitative method that can report on the time evolution of distance distributions between labeled residues. The contrast between lit and dark states will be enhanced by maximizing the fraction that are loaded with a chromophore (further discussion of lineshape interpretation in S.I. 2.5). The effects of rotational averaging may be mitigated by incorporating proteins into stabilizing agents (*e.g*., hydrogels) and enabling the extraction of time-dependent distance distributions from TiGGER will require recording the entire field-swept cwEPR spectrum as a function of time rather than the cwEPR signal at a fixed magnetic; rapidscan EPR offers an attractive path towards this goal. Additionally, methods that have been developed to extract distance distributions from nitroxide lineshapes will need to be extended to Gd(III) spin labels at high fields (e.g., [77, 78]).

## 4 Conclusion

In this paper, we demonstrated the first step to “filming” a protein in action using TiGGER. Site-specific labeling allowed us to track a change in distances between a pair of residues in an ensemble of room temperature proteins in solution state. We confirmed a mutation-induced slowing of the mechanical photocycle, highlighting the importance of TiGGER to the optogenetic community. Additional experimental development and an application of TiGGER to a range of protein residues will enable three-dimensional, time-dependent mapping of protein mechanical action, and may play an important role in improving design of optogenetic actuators and fluorescent reporters.

The technique presented here should be applicable to studying conformational changes triggered by other factors as well, such as ligand binding by rapid mixing, voltage actuation, or temperature jumps. We hope in the future that TiGGER may be used to complete stories of protein triggered functional dynamics that are partially told by time-resolved X-ray crystallography and IR spectroscopy, FRET, freeze-quench cryo-EM, solid-state NMR, and DEER, among others.

## 5 Acknowledgements

The authors would like to thank Karen Tsay for her thoughtful insight and assistance running Q-band DEER experiments. We would also like to acknowledge support from the NSF though grant MCB 2025860 and the UC Ofce of the President Multicampus Research Programs and Initiatives under MRI-19-601107 for the development of TiGGER and the core research presented here. Biochemical and complementary biophysical studies of AsLOV2 were supported by the NIH MIRA through grant R35GM136411. JEL thanks The Royal Society for a University Research Fellowship. Figures were created in Python using SciencePlots.[79] All data and programs used for this publication are available at [80] and [81], respectively.

