## Supplemental Information for "Triggered functional dynamics of AsLOV2 by time-resolved electron paramagnetic resonance at high magnetic fields"

#### 1 Experimental procedures

##### 1.1 Protein expression

A DNA fragment encoding the LOV2 domain of Avena sativa phototropin 2 (residues 403-546) cloned into pET-26b (+) vector via NdeI (5') and XhoI (3') cloning sites with a C- terminal 6X histidine affinity tag was purchased from GENE UNIVERSAL INC. The plasmid was transformed into E. coli BL21 (DE3) cells and successful transformants were grown in 10 mL LB media containing kanamycin (5 $\mu$g/mL) at 37 °C under continuous shaking for 18 hours. After overnight shaking, the culture is then diluted into a 1L LB media containing kanamycin and grown at 37 °C under continuous shaking. At an OD600 of 0.6-0.8, the culture was induced with 1mM IPTG and incubated for 16 hours at 18 °C in the dark. The cell culture was then harvested by centrifugation at 5000 g for 20 min and resuspended in 30 mL lysis buffer (20 mM TrisHCl, 500 mM NaCl pH: 8.00) along with 5 mM $\beta$-mercaptoethanol and lysozyme (1 mg/mL). After an incubation of 30 mins at 4 °C, the resuspended cell culture was lysed with sonication (60 cycles, 10 seconds pulses with an interval of 10 seconds) and centrifuged at 11000 rpm for 30 mins at 4 °C to remove cell debris.

##### 1.2 Protein purification

In order to reduce the amount of apo-protein, exogeneous flavin mononucleotide was added to the protein in excess (150X). The mixture is incubated at room temperature in the dark under rotation (15 rpm) for 30 mins. For purification, the mixture is further incubated with 4 mL of Ni-NTA slurry and kept under rotation (15 rpm) at 4 °C in the dark for 2 hours. The mixture is then allowed to rest so that it can separate into two layers. The supernatant is then carefully removed and the sediment with the required protein bound Ni-NTA is washed 3 times each with 40 mL wash buffer (20 mM TrisHCl, 500 mM NaCl, 40 mM imidazole, 5 mM $\beta$-mercaptoethanol, pH: 8.00). For elution, the Ni-NTA bound protein slurry is loaded into a gravity flow chromatographic column and eluted with 10 mL elution buffer (20 mM TrisHCl, 500 mM NaCl, 500 mM imidazole, 5 mM $\beta$-mercaptoethanol, pH: 8.00). The eluted protein was then stored at 4 °C overnight. The stored protein is then further purified using a size exclusion chromatographic column, where AsLOV2 was run through a HiLoadTM 16/600 SuperdexTM 200 pg column (GE Healthcare, Chicago, IL) connected to a NGC$^{TM}$ Medium-Pressure Liquid Chromatography System (BioRad, Herculus, CA) with the elution buffer (20 mM TrisHCl, 150 mM NaCl, pH: 8.00). The eluted protein is concentrated and incubated with 10 mM DTT under rotation at 4 °C overnight to reduce the disulfide bonds in order to spin label the following day.

##### 1.3 Protein spin labeling

For labeling AsLOV2 with the nitroxide spin label (MTSL) (1-Oxyl-2,2,5,5-tetramethyl-Δ3-pyrroline-3-methyl methanesulfonothioate) as well as Gd-sTPATCN, purified AsLOV2 was first treated with 10 mM DTT at 4 °C overnight to completely reduce the thiol groups on cysteines. After removing DTT with a desalting column (PD-10, GE Healthcare, Chicago, IL) , AsLOV2 was immediately mixed with 10-fold molar excess of the spin labels and the mixture was gently shaken overnight at room temperature. Excess spin label was removed by size exclusion chromatography (HiLoadTM 16/600 SuperdexTM 200 pg column GE Healthcare, Chicago, IL) using an NGC$^{TM}$ Medium-Pressure Liquid Chromatography System (BioRad, Herculus, CA). The spin-labeled AsLOV2 is concentrated to about 1.5 mM protein prior to use for hfEPR measurements and to about 30-40 µM protein for UV-Vis measurements.

#### 1.4 Protein Enrichment

Because TiGGER is based on a cwEPR lineshape measurement, signal from singly-labeled proteins forms an unwanted background that can obscure the desired signal from doubly-labeled proteins. In order to enrich the concentration of doubly-labeled protein, the spin-labeled preparation was further incubated at room temperature for 50 min. with 0.01 mg/mL biotin-maleimide (BM) stock solution (prepared in 20 mM Tris-HCl 150 mM NaCl pH: 8.00) to react any residual free (*i.e.*, unlabeled) cysteines in AsLOV2 with BM (green-blue stars in main text Fig. 2, top). BM was added to achieve a 1:1 molar ratio between AsLOV2 cysteines and the BM. After incubation, the mixture was added to a streptavidin-agarose resin (Pierce®, Thermo Scientific, main text Fig. 2, right) to bind with biotinylated AsLOV2 thereby allowing DL AsLOV2 proteins to be enriched in the flow-through.

As per section 1.2 of the main text, protein enrichment also had the intended effect of increasing dipolar broadening. The increased broadening is evident in 87 K cwEPR field-swept experiments of enriched and unenriched doubly Gd-labeled AsLOV2 shown in Fig. 1.

This is the first report of this enrichment technique, which was invented by JEL.

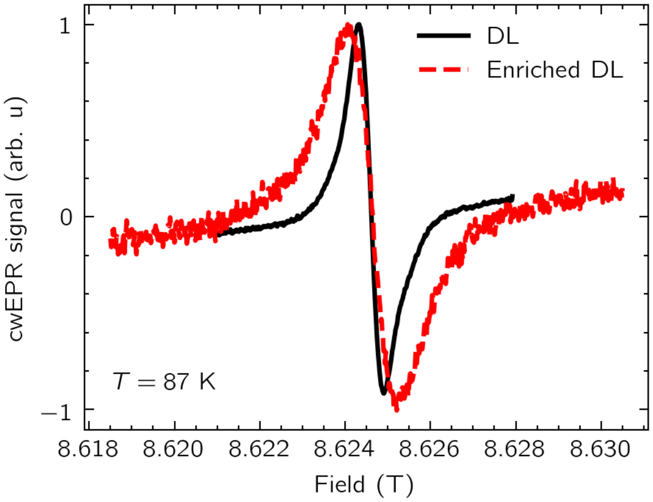


Figure 1: Effect of enrichment on dipolar broadening. Broadening was more significant after enrichment due to a larger fraction of spins being dipolar coupled (dashed red line, enriched; solid black line, unenriched).

The spectral changes observed in the room temperature hfEPR of Q513A DL and C450 DL were smaller than those of DL. To ensure that this was not due to reduced double labeling yield, we compared their lineshapes at 87 K. Spectra shown in Fig. 2 show that the linewidths are indistinguishable and therefore that double label yield and interspin distance must be very similar for all three.

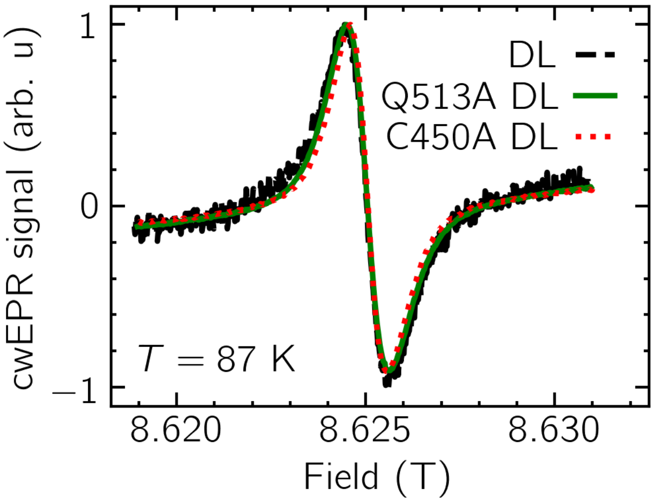


Figure 2: Comparison of wildtype, C450A, and Q513A lineshapes at 87 K. There was no significant change in linewidth due to the mutations. This means that double-labeling efficiency was similar for all three samples and that the restricted motion we saw for C450A and Q513A was due to an inhibited photocycle, not insufficient labeling. Signal-to-noise varies due to varying sample concentration.

##### 1.5 Q-band DEER

As a test of the effectiveness of the enrichment procedure, we measured DEER spectra of unenriched and enriched samples, as well as of a 2.8 nm MTSL ruler that we assumed to have 100% labeling. DEER measurements were performed at 65 K with 11 ns pump pulse at Q-band (34 GHz Bruker E580 ELEXSYS pulse EPR spectrometer equipped with a TWT amplifier at 300 W). The purified enriched doubly-MTSL-labeled AsLOV2 gave a modulation depth of ∼0.4, as shown in Fig. 3(compared to a modulation depth of ∼0.35 for un-enriched DL AsLOV2). The 2.8 nm MTSL ruler also gave a modulation depth of ∼0.4, as shown in Fig. 4. These results indicate that we achieved close to 100% double labeling efficiency after enrichment.

Effects of light activation were also measured using DEER. The large peak in Fig. 3 just above 2 nm decreases by about 25% upon light activation, and the tail at distances between 2.8 and 6 nm increases correspondingly. Thus, even with a high double-label yield, the number of intra-protein spin pairs that measurably moved away from each other is small. This highlights the sensitivity of TiGGER: even from a small fraction of spins moving apart, the signal from TiGGER was clear and significant.

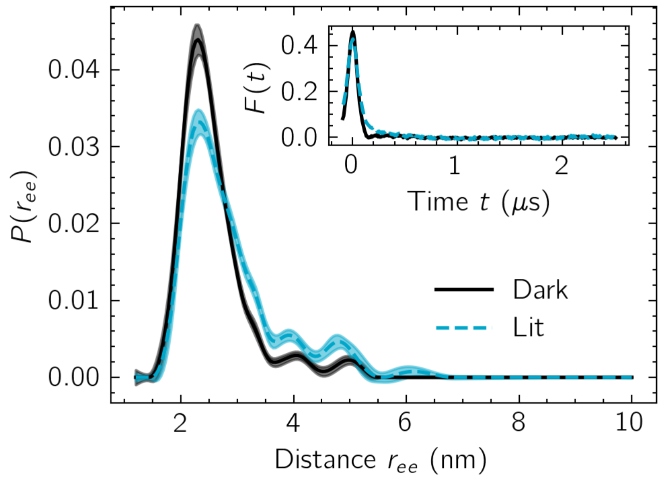


Figure 3: Effect of light-activation on inter-residue spacing of AsLOV2. Q-band DEER data (with 11 ns pump pulse) of doubly-MTSL-labeled AsLOV2 at the sites T406C and E537C in the dark (solid black) and the lit (dashed blue) state. Background-corrected dipolar evolutions of the dark and the lit state (inset) and their corresponding distance probability distributions from Tikhonov regularization with LongDistances1020 (June 29, 2021 version, [1]) are shown.

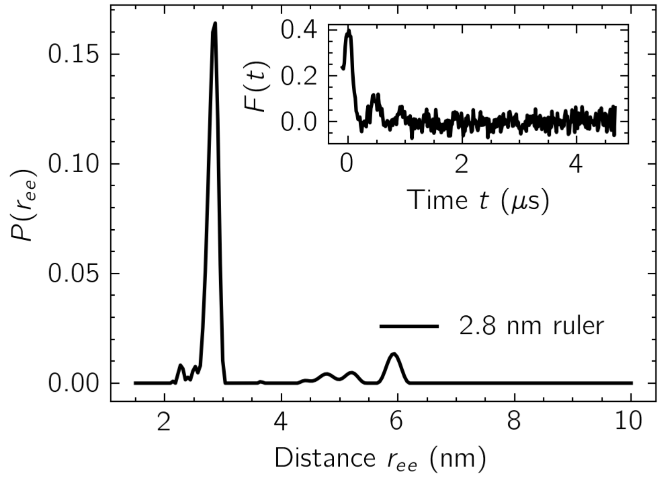


Figure 4: Q-band DEER data (with 11 ns pump pulse) of a 2.8 nm nitroxide ruler. Background-corrected dipolar evolutions of the 2.8 nm ruler (inset) and corresponding distance probability distributions from Tikhonov regularization with LongDistances1020 (June 29, 2021 version, [1]) are shown.

#### 1.6 Effect of SDSL on protein dynamics

As shown in Figs. 5-8, site-directed spin labeling had no effect on the UV-Vis spectral features or lifetimes of AsLOV2 mutants (compare DL AsLOV2, Fig. 5 to unlabeled AsLOV2, main text Fig. 1, middle left; compare DL Q513A AsLOV2, Fig. 7 to unlabeled Q513X AsLOV2 in [2]). Note the shift in peak absorption between DL T406C-E537C and DL T406C-E537C Q513A indicates a change in electronic environment near the FMN due to mutation of Q513 to alanine, as is reported elsewhere.[2–4]

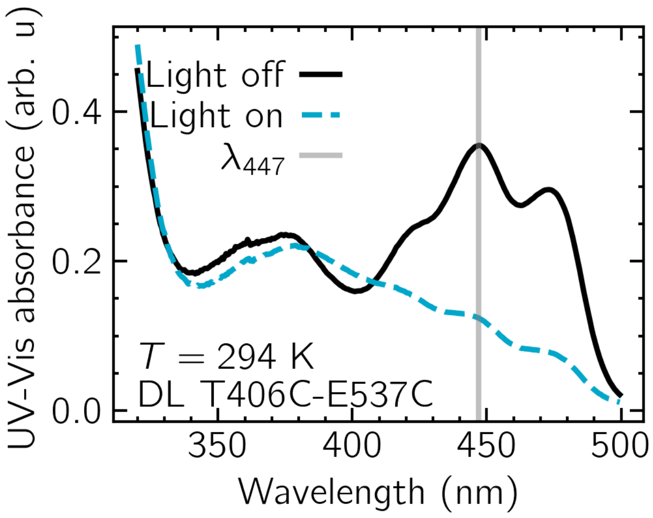


Figure 5: UV-Vis absorption spectra of AsLOV2 T406C-E537C with (dashed blue line) and without (solid black line) blue light activation (Thorlabs, Inc. LIU470A). The gray line indicates the wavelength at which the lifetime of the protein was measured.

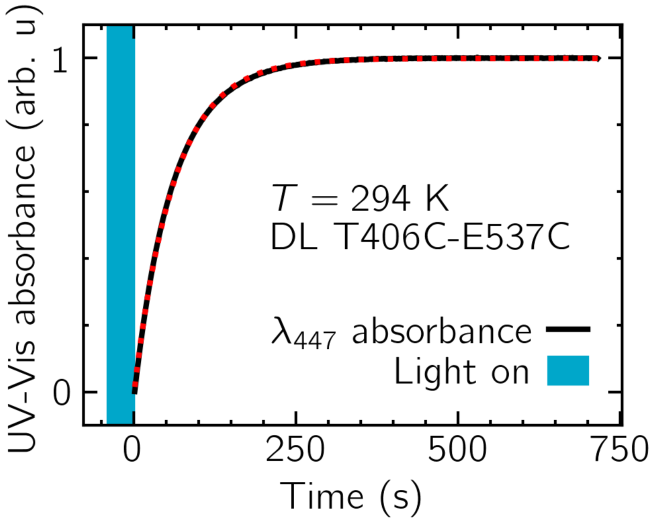


Figure 6: The lifetime of the protein ($\tau = 61.91 \pm 0.07$ s) after activation with blue light was measured by recording the UV-Vis absorbance at 447 nm.

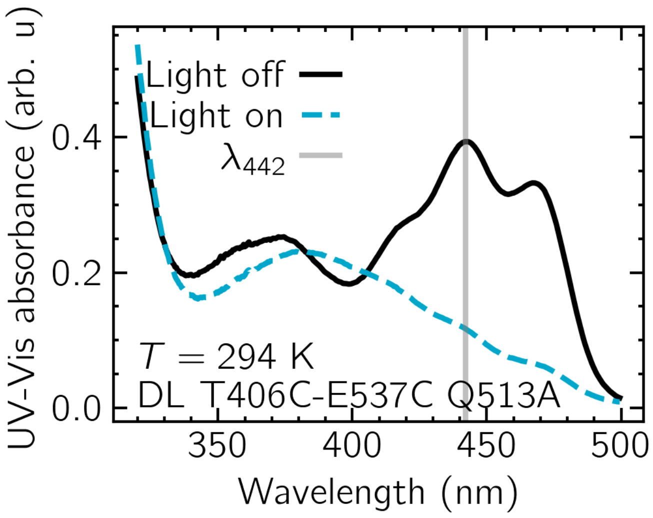


Figure 7: UV-Vis absorption spectra of AsLOV2 Q513A T406C-E537C with (dashed blue line) and without (solid black line) blue light activation (Thorlabs, Inc. LIU470A). The gray line indicates the wavelength at which the lifetime of the protein was measured.

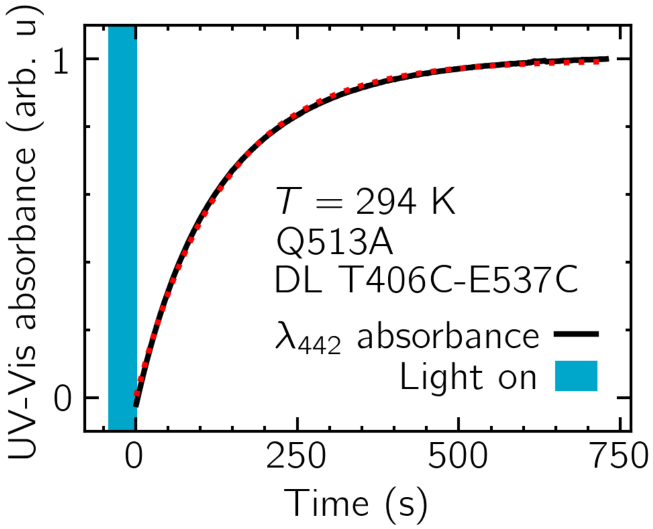


Figure 8: The lifetime of the protein ($\tau = 135.4 \pm 0.3$ s) after activation with blue light was measured by recording the UV-Vis absorbance at 442 nm.

#### 1.7 Additional protein sample details

Though it was not expected to have any meaningful effects on our results, some erroneous labeling of the central cysteine did occur during the labeling process, as shown in Figure 9. The central cysteine is critical for photo-absorption, however, and therefore any erroneously labeled C450 would neither be light-activated nor affect the results. The spin labeling chemistry involves reaction of the spin label to a thiol group (S-S for MTSL and C-S for Gd-sTPATCN), so when C450 is spin labeled it becomes unavailable to form the disulfide bond with the FMN (required for forming the photo-adduct). We confirmed this as we observed no photoactivity after intentionally mutating the central cysteine to alanine (C450A) to simulate the effect of labeling (Fig. 10).

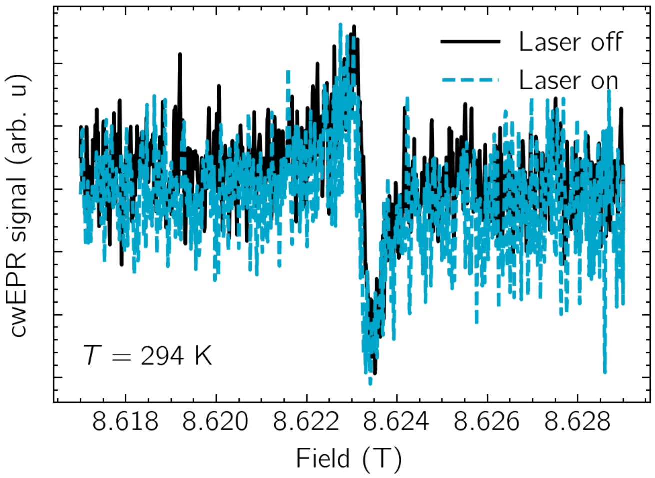


Figure 9: cwEPR spectra of wildtype AsLOV2 after undergoing labeling procedure. Since there are no cysteine mutations, sites 406 and 537 are unlabeled and all EPR signal is due to coincidental labeling of the central cysteine. The frequency of this occurrence is estimated at $\sim 1\%$ by 240 GHz and Q-band EPR.

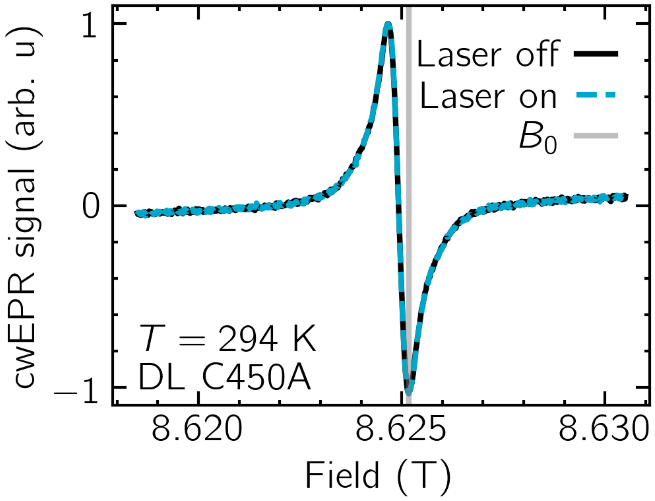


Figure 10: cwEPR spectra of C450A AsLOV2 after undergoing labeling procedure. Since the central cysteine is mutated, the protein does not photoswitch and no lineshape change is absorbed between laser off (solid black line) and laser on (dashed blue line).

#### 1.8 EPR spectrometer

High-field EPR was carried out at UCSB's Institute for Terahertz Science and Technology (ITST). ITST's homebuilt EPR spectrometer consists of a 12.5 T field-swept magnet (Oxford Instruments), 60 mW 240 GHz cw source, and subharmonically mixed heterodyne receiver (Virginia Diodes, Inc., Charlottesville, VA) that operates in induction mode and has been described in detail previously.[5, 6] A small offset of the recorded magnet field of approximately 12 mT causes Gd(III) resonances shown to appear at 8.62 T, not 8.608 T as would be expected for Gd labels with an isotropic $g$-value of 1.992.[7] A 240 GHz Gaussian beam was coupled by a corrugated waveguide into the sample space, where liquid samples were loaded into a 100 $\mu$m-thick, 2-by-5 mm borosilicate glass capillary (VitroCom, Mountain Lakes, NJ) to maximize the ratio of optical surface area to optical density.[8] Capillaries were then placed on a Teflon® tape-covered, 7 mm wide, protected silver mirror. Field-swept EPR experiments were done at first in the dark and then under 450 nm laser illumination. The laser produced 70 mW at 450 nm (Laser Components USA, Inc., Bedford, NH) and was coupled into a fiber optic that carried approximately 15 mW to the sample space. Time-dependent experiments were completed by continuously collecting field-modulated, lock-in detected, cwEPR data as a function of time (60 ms time steps) and activating the laser light for 10 cycles of 5 seconds on, 175 seconds off. The repetitions were then averaged to reduce noise fluctuations (see S.I. section 2.1).

### 2 Results and discussion

#### 2.1 Transient EPR

To record transient trEPR data, the field was fixed while field-modulated, lock-in detected data was collected. Each experiment was repeated ten times continuously at fixed intervals and were averaged in post-processing. The results of this process is shown for singly-labeled proteins in Figs. 11 and 12 (small trEPR signal), for doubly-labeled protein in Fig. 13 (much larger trEPR signal), and doubly-labeled protein with the central cysteine mutated out in Fig. 14 (no detectable trEPR signal).

There was no appreciable effect of repeated illumination on the sample (for example due to bleaching), as shown by Figure 15. Fig. 15 shows fit amplitude and decay time for each individual run and no obvious pattern with increasing experiment number was observed.

#### 2.2 Dipolar Hamiltonian

The high field approximation allows us to express the dipolar contribution to the spin Hamiltonian as

$$\mathcal{H}_{dd} = \omega_{dd}^0 \left( S_z^A S_z^B - \frac{1}{4} \left( S_+^A S_-^B + S_-^A S_+^B \right) \right) \cdot \left( 3\cos^2\theta_d - 1 \right) \tag{1}$$

where $S_z$ is the standard spin-7/2 spin operator, $S_\pm$ are the non-Hermitian raising and lowering operators, $\omega_{dd}^0 = \frac{\mu_0}{4\pi}\frac{\mu_b^2 g_1 g_2}{\hbar}\frac{1}{|\vec{r}_{AB}|^3}$ is the dipolar splitting caused by one spin on another and $\theta_d$ is the angle between $\vec{r}_{AB}$ and the applied field $B_0$.[9, 10] Increasing the ensemble-averaged spin-spin separation, $|\vec{r}_{AB}|$, reduces the dipolar contribution to the spectrum and thereby narrows the cwEPR line. The distance dependence of the dipolar coupling is commonly applied in DEER to measure static inter-spin distance, but may be extended to cwEPR for real-time tracking of ensemble-averaged spin-spin separation, assuming, for example, that $\mathcal{H}_{dd} \gg \mathcal{H}_{\text{ZFS}}, \mathcal{H}_{\text{hyperfine}}$, etc. An important note about equation (1) is that it will average to zero with fast tumbling where $\omega_{dd}\tau_c \ll 1$, where $\tau_c$ is the rotational correlation time.[11] If persistent dipolar broadening of doubly-labeled proteins in solution is observed, it indicates that $\omega_{dd}\tau_c$ must be smaller than or comparable to 1 under our experimental condition.

#### 2.3 scipy.optimize.curve_fit() fitting

The range of values for $\tau$ throughout the main text and SI give a 95% confidence interval for the fit parameters; this was calculated by doubling the standard deviation for each parameter as given by the covariance matrix from curve_fit().

#### 2.4 Single-label relaxation

Though small compared to the DL T406-E537C sample, it is worth noting that we see an unexpected nonzero temporal decay of the singly-labeled samples. These small lineshape changes may come from a small percentage of spins experiencing unintended dipolar broadening and subsequent narrowing as a result of non-specific labeling. Alternatively, there might be a small contribution from inter-AsLOV2 interaction that give rise to dipolar coupling between two singly spin labeled AsLOV2 that, upon light activation, move further apart from each other.

#### 2.5 Lineshape interpretation

A clear observation of dipolar broadening of Gd-sTPATCN requires that the distance between labels is smaller than about 4 nm. In the dark state, two protein configurations – singly-labeled proteins, and dark-state-unfolded proteins – could have Gd-Gd distances much longer than 4 nm, and hence give a lineshape that is not dipolar broadened. In the dark, the cwEPR spectrum is thus a superposition of a sharp line, associated with singly-labeled and dark-state unfolded proteins, and a dipolar-broadened line, associated with doubly-labeled, dark-state folded proteins. It is likely that even after purification, a few percent of the proteins are still singly labeled. Further, it is known that not all proteins will activate upon illumination, as a small fraction have a labeled central cysteine (C450), others are empty (without FMN chromophore), and some are already unfolded.[12] Upon light activation, only the lineshapes of doubly-labeled, dark-state folded proteins with a FMN chromophore will unfold. Other populations contribute to background obfuscation. Hence, there is more work to do how to delineate the action of the AsLOV2 population of interest.

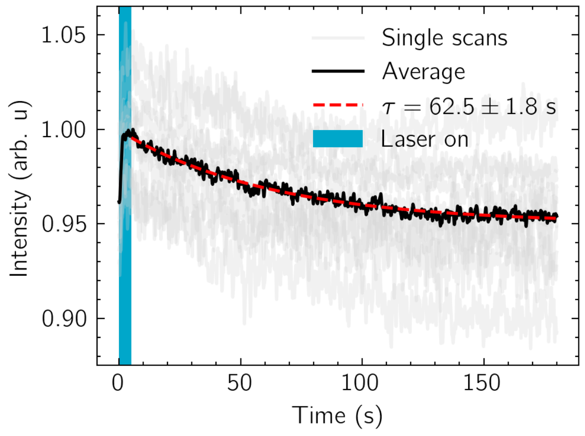


Figure 11: Time-dependent cwEPR signal change of T406C AsLOV2 at $B_0$ (vertical gray line in main text figure) showing increase during the laser pulse (solid blue vertical line) and subsequent decay after the laser is turned off. Amplitude of singly labeled decay is much smaller than that of doubly labeled. Gray solid lines, black solid line, and red dashed line correspond to single scans, their average, and exponential fit to the decay, respectively.

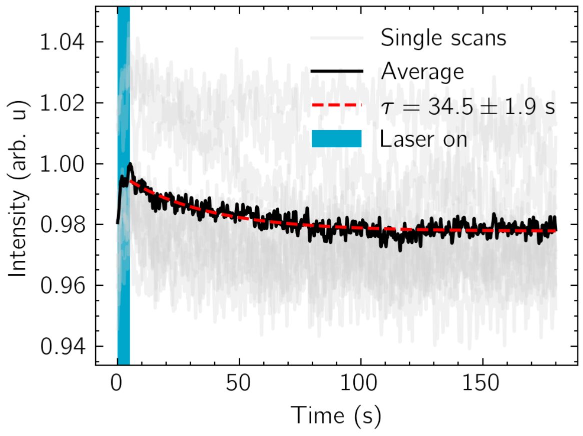


Figure 12: Time-dependent cwEPR signal change of E537C AsLOV2 at $B_0$ (vertical gray line in main text figure) showing increase during the laser pulse (solid blue vertical line) and subsequent decay after the laser is turned off. Amplitude of singly labeled decay is much smaller than that of doubly labeled. Gray solid lines, black solid line, and red dashed line correspond to single scans, their average, and exponential fit to the decay, respectively.

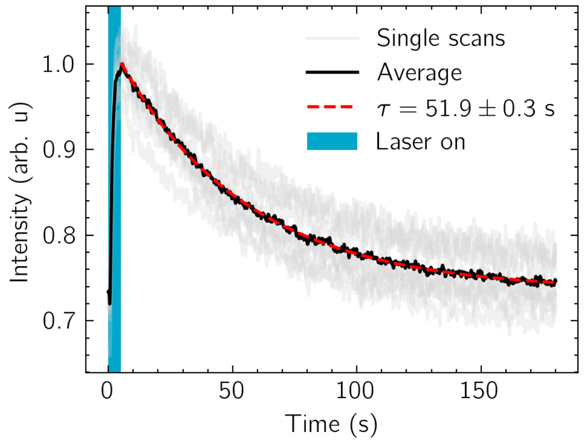


Figure 13: Time-dependent cwEPR signal change of T406C-E537C AsLOV2 at $B_0$ (vertical gray line in main text figure) showing increase during the laser pulse (solid blue vertical line) and subsequent decay after the laser is turned off. Gray solid lines, black solid line, and red dashed line correspond to single scans, their average, and exponential fit to the decay, respectively.

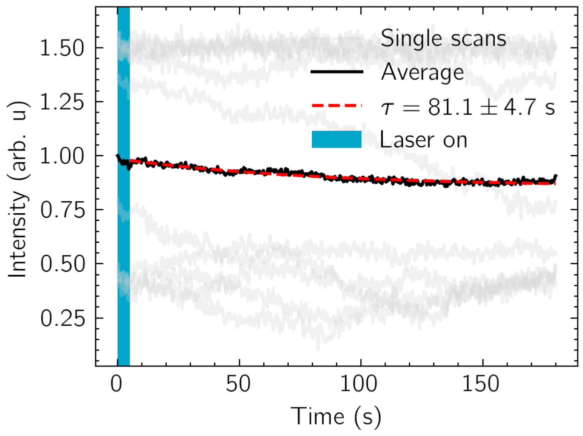


Figure 14: Time-dependent cwEPR signal change of C450A T406C-E537C AsLOV2 at $B_0$ (vertical gray line in Fig. 10) showing lack of change during laser pulse (solid blue vertical line) and afterward. Gray solid lines, black solid line, and red dashed line correspond to single scans, their average, and exponential fit to the decay, respectively. Single scan lines do not overlay well because of a small amount of spectrometer drift and a complete lack of signal; removing baseline and normalizing to maximum creates seemingly random peak intensities.

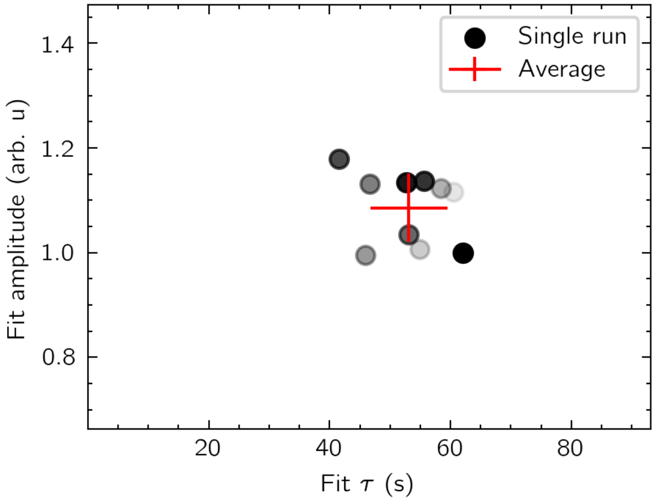


Figure 15: Fit parameters for each individual experiment on DL T406C-E537C AsLOV2. Filled in gray circles represent one experiment, lighter color means individual experiment happened later. Tight clustering, a lack of discernible pattern, and small standard deviation (represented by error bars in red) demonstrate no discernible change to the sample during subsequent runs.

### 3 Author contributions

S. Maity: methodology, validation, investigation, writing original draft, writing review and editing, visualization.
B. Price: methodology, validation, investigation, writing original draft, writing review and editing, visualization, formal analysis, software.
C. B. Wilson: conceptualization, methodology, writing review and editing.
A. Mukherjee and M. Z. Wilson: conceptualization, resources, writing review and editing.
M. Starck and D. Parker: resources (Gd-sTPATCN spin label synthesis).
J. E. Lovett: conceptualization, resources, methodology, writing review and editing.
S. Han and M. S. Sherwin: conceptualization, methodology, writing original draft, writing review and editing, supervision, project administration, funding acquisition.
